# Compound heterozygous *ZP1* mutations cause empty follicle syndrome in infertile sisters

**DOI:** 10.1101/363911

**Authors:** Ling Sun, Xiang Fang, Zhi-Heng Chen, Han-Wang Zhang, Xiao-Fang Peng, Yu Deng, Ting Xue, Min-Na Yin, Qian-Ying Zhu, Chun-Lin Liu, Na Li

## Abstract

**Purpose:** Empty follicle syndrome (EFS) is a condition in which no oocyte is retrieved from mature follicles after proper ovarian stimulation in an *in vitro* fertilization (IVF) procedure. Genetic evidence accumulates for the etiology of recurrent EFS even with improved medical treatment which had avoided the pharmacological or iatrogenic problems. Here, this study investigated the genetic cause of recurrent EFS in a family with two infertile sisters.

**Methods:** In this work, we present two infertile sisters in a family with recurrent EFS after three cycles of standard ovarian stimulation with hCG and/or GnRHa therapy. We performed whole-exome sequencing and targeted sequencing in the core members of this family, and further bioinformatics analysis to identify pathogenesis of gene.

**Results:** We identified compound heterozygous variants, c.161_165del (p.54fs) and c.1166_1173del (p.389fs), on zona pellucida glycoprotein 1 (*ZP1*) gene, which were shared with two infertile sisters. Cosegregation tests on the affected and unaffected members of this family confirmed that the allelic mutants were transmitted from either parent.

**Conclusions:** This EFS phenotype was distinct from the previously reported disruption of zona pellucida due to homozygous *ZP1* defects. We thus propose that the specific mutations in *ZP1* gene may render a causality for the recurrent EFS.

## Introduction

During in vitro fertilization (IVF) treatment, multiple follicles developed following ovarian stimulation. After trigger with human chorionic gonadotrophin (HCG), cumulus-oocyte complexes (COCs), which consist of cumulus cells surrounding the centrally located oocyte, can be isolated from the follicular fluid. However, in rarely condition, no identifiable COCs were collected; this phenomenon is defined as empty follicle syndrome (EFS)[1].

EFS can be classified as either “false” EFS or “genuine” EFS. The “false” EFS was due to the inappropriate timing and dosage of HCG administration[2–4], after rescue protocols[5–10], the oocytes could be harvested; whereas “genuine” EFS was termed as without oocytes nor cumulus-corona complexes being retrieved after improved protocols[11–13], and it was implied underlying genetic defects.

Studies identified homozygous mutations in the gene of *LHCGR* (MIM#152790) in EFS [14, 15]. *LHCGR* gene encodes the luteinizing hormone/choriogonadotropin receptor; the impaired functioning of LHCGR caused diminished response to the administration of beta-hCG. More recently, based on genome-wide linkage analyses and whole-exome sequencing, a study revealed a recurrent heterozygous missense mutation in zona pellucida glycoprotein 3 (ZP3, MIM#182889) in two EFS families [16]. The author proposed that via dominant negative inhibition, this mutation could destroy the assembly of the zona pellucid (ZP) thus leading to oocyte degeneration and EFS. However, the etiology research on EFS, especially those recurrent ones, is far from completeness, therefore, the full spectrum of genes and mutations involved in EFS are still to be expanded.

In this study, we identified compound heterozygous mutations of *ZP1* gene (MIM#195000) from a family with two sisters of abnormal oogenesis with EFS. The allelic mutants were transmitted from either parent. We hypothesized that these two ZP1 mutations hurdled the oocyte maturation at an early stage, thus incurring degeneration and empty follicles.

## Results

### Clinical description

The proband was enrolled in IVF treatment due to the asthenospermia and teratozoospermia of her husband (Table S2). In the first attempt, a gonadotropin-releasing hormone (GnRH) agonist protocol was performed (Table S3). On the day of trigger, the peak estradiol was 5757 pmol/ml. Five follicles were documented by ultrasound, measuring 18–20 mm average diameter, and totally twelve follicles were recorded. Oocyte retrieval was performed 36 hours after recombinant human chorionic gonadotropin (r-hCG,Ovidrel^®^,Merck Serono, Germany) administration. No recognizable oocytes were retrieved after thorough aspiration and flushing of the seven follicles in the right ovary, so the oocyte retrieval was suspended and the HCG level in blood was evaluated immediately. Because of the normal hormones assays, ovum pick up (OPU) was performed 2 hours later, and five follicles in the left ovary were aspirated. Several cumulus–corona complexes were recovered but no recognizable oocytes inside. The second IVF attempt was carried out with GnRH antagonist protocol (Table S3) 3 months later; dual trigger was administered. However, similar to the first attempt, no recognizable oocytes were retrieved.

In case 2, after GnRH antagonist treatment, nine cumulus–corona complexes were collected (Figure 1B), however, no recognizable oocytes were found as well. After removal of the cumulus cells, seven degenerated oocyte cytoplasts were identified, without zona pellucida surrounding, among which five had germinal vesicle (GV) body (Figure 1D). In contrast, the cumulus cells of controls were radiating and well expanded (Figure 1A), and after removal of the cumulus cells, the control oocyte exhibited a clear cytoplasm with homogeneous fine granularity, surrounded with insoluble zona pellucida (Figure 1C).

**Figure 1.**
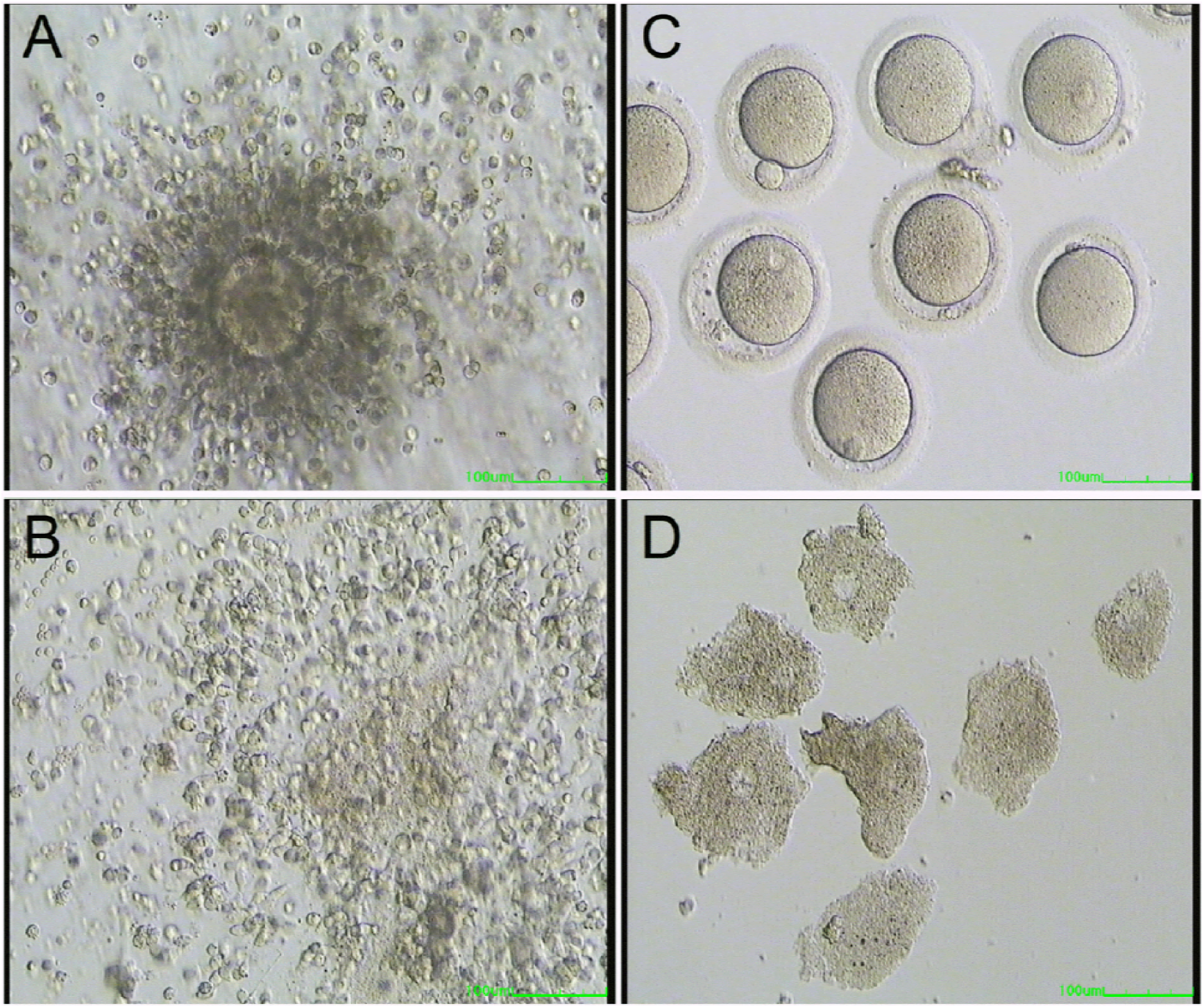
Phenotypic features of the proband. (A) shows a normal oocyte with cumulus-corona complexes. (B) shows cumulus-corona complexes of the proband. (C) shows the control oocytes after removing the cumulus cells. The control oocyte exhibits a clear cytoplasm with homogeneous fine granularity, surrounded with insoluble zona pellucida. (D) shows the degenerated oocytes after removing the cumulus cell from the proband. Under GnRH antagonist protocol, nine cumulus–corona complexes were collected, however, no recognizable oocytes were found. After removal of the cumulus cells, seven degenerated oocyte cytoplasts were identified, without zona pellucida surrounding, among which five had germinal vesicle (GV) body. Scale bar: 100 um.

### Compound heterozygous mutations in ZP1

We performed Whole-exome sequencing (WES) on the proband of the family (Figure 2A II-3). Then followed by filtering, annotation, and prioritization for the WES data on the proband, the cosegregation test was performed for the selected candidate variants on all six family members via Sanger sequencing. This strategy revealed a compound heterozygous variants, c.161_165del (p.54fs) and c.1166_1173del (p.389fs) on *ZP1* (transcript: NM_207341), as a potential pathogenic variant (Figures 2B and 2C). These compound heterozygous variants were shared by the proband and the affected sister (Figure 2A II-2). Cosegregation tests on the affected and unaffected members of this family confirmed these mutations were transmitted with the manifestation when in compound heterozygotes (Figure 2A and 2C).

**Figure 2.**
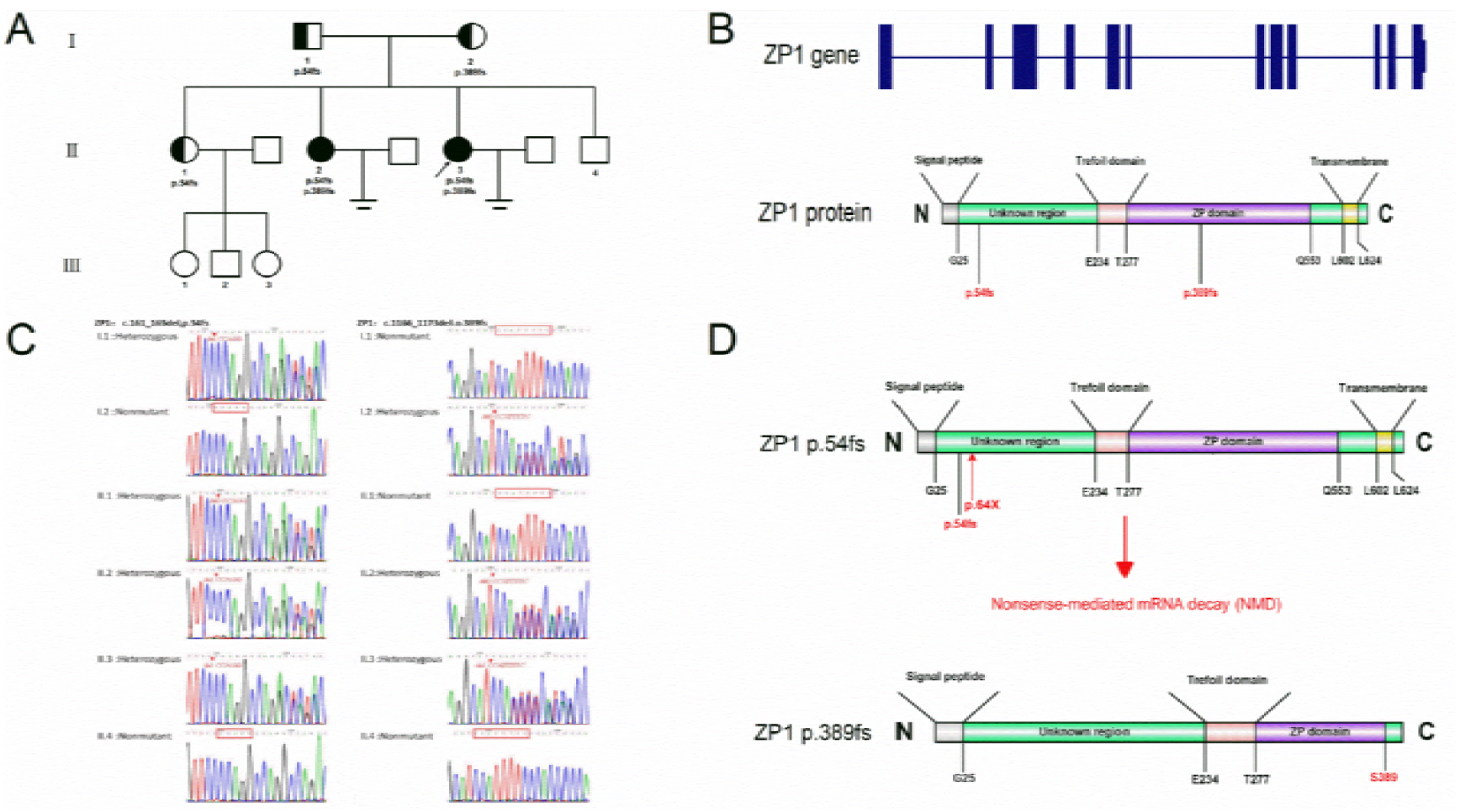
Details of pedigree, gene, mutations, and Sanger-sequencing confirmation. (A) Pedigree information. Squares indicate male family members, and circles indicate female family members. Solid symbols represent affected subjects, half solid denoting the unaffected mutation carriers, white indicating unaffected subjects. Equal signs infertility. The arrow points to the proband of the family. The specific mutations of the zona pellucida 1 (*ZP1*) are indicated for the affected family members. (B) The schematic diagram of ZP1 with functional domains and the location of the identified mutations. ZP1 peptide contains an N-terminal signal sequence (gray), three unknown regions (green), a Trefoil domain (pink), a ZP domain (purple) and a transmembrane domain (yellow). The individual amino acids plus positions indicate the domain boundaries. The locations of the identified mutations are showed in red. (C) Sanger sequencing chromatograms of allelic mutations of *ZP1* in all family members. Two family members (I-1 and II-1) carry a heterozygous frameshift deletion of 5 nucleotides (nt), c.161_165del in *ZP1.* This mutation leads to the formation of a premature stop codon (p.54fs64X), in exon 1 of *ZP1*, which is predicted to cause the nonsense-mediated mRNA decay (NMD). Family I-2 carries a frameshift deletion of 8 nucleotides (nt), c.1166_1173del (p.389fs), in exon 7 of ZP1. This mutation, causes a premature stop codon (I390fs404X), results in a truncated protein of 404 amino acids, instead of 638 amino acids of the wildtype ZP1 protein. The proband (II-3) and the affected sister (II-2) have both of these two frameshift deletions. Family II-4 shows the wildtype genotype. (D) The schematic diagram of allelic mutations of ZP1 protein. The frameshift deletion, i.e., c.161_165del (p.54fs), leads to the formation of a premature stop codon (p.54fs64X), which is predicted to cause NMD. The other frameshift deletion, i.e., c.1166_1173del (p.389fs), causes a truncated protein by a premature stop codon (I390fs404X), which consists of the N-terminal signal sequence, the trefoil domain, and the first half of the zona pellucid domain. The other domains, such as the transmembrane domain, part of the zona pellucida domain, and other C-terminal domains, are probably absent.

The candidate gene was *ZP1* (MIM#19500), a well-known gene encoding one of four zona pellucida glycoproteins [17], namely, *ZP1*, *ZP2* (MIM#182888), *ZP3* (MIM#182889), and *ZP4* (MIM#613514). It has been shown that ZP1 was a critical component for the structural integrity of the zona matrix in mice models [18].

Notably, the heterozygous frameshift deletion of 5 nucleotides (nt), c.161_165del (p.54fs), which we found in exon 1 of *ZP1* (transcript: NM_207341), was transmitted from the father of the proband (Figure 2A I-1) and also carried by the unaffected sister (Figure 2A II-1), who had already given birth for three times (Figure 2C). This mutation led to the formation of a premature stop codon (p.54fs64X), in exon 1 of *ZP1*, that was predicted to cause the nonsense mediated mRNA decay (NMD)[19] (Figure 2D), indicating that there would be no corresponding mature mRNA generated. This mutation was absent in our in-house ethnic-matched 2,200 individual genomes and the public databases including 1000Genome, ExAc database and ESP6500 database. Interestingly, the second heterozygous mutation, a frameshift deletion of 8 nucleotides (nt), c.1166_1173del (p.389fs), in exon 7 of *ZP1* (transcript: NM_207341), transmitted from the mother of the proband (Figure 2A I-2) (Figure 2C), was the same mutation reported by a previous work[20]. In that work, Huang et al. found in a familial infertility case a homozygous mutation, caused a premature stop codon (I390fs404X), which resulted in a truncated protein of 404 amino acids, instead of the 638-amino-acid original ZP1 protein (Figure 2D) and posited that the defective ZP1 hindered the transportation of ZP3, another key elements in zona pellucida, from the cytoplasm to the surface of oocyte[20]. Thus in our case, since one allelic mutation caused NMD and the other truncated the functional protein, these compound heterozygous mutations of *ZP1* genes would presumably lead to half-dosage truncated ZP1 of the previous reported case and worsen the pathological condition, so may well be responsible for the specific phenotype of EFS.

## Material and Methods

This study was approved by the Reproductive Medical Ethics Committee of Guangzhou Women and Children’s Hospital.

### Case I

The proband of the family (Figure 2A II-3) was a 23-year-old woman with a 3-year history of primary infertility. She had regular menses (26–28 days in length) since menarche at the age of 16. Her ovarian reserves were generally normal with a total of 12 antral follicles revealed by ultrasound and normal basal sex hormone and anti-Müllerian hormone levels (Table S1). The karyotyping result of proband was 46, XX, with no observable chromosomal abnormality. The proband was enrolled in IVF treatment due to the asthenospermia and teratozoospermia of her husband (Table S2). In the first attempt, a gonadotropin-releasing hormone (GnRH) agonist protocol was performed (Table S3); after down-regulation, 150 IU of recombinant FSH (Gonal-f^®^,Merck Serono, Germany) was administered for 10 days. On the day of trigger, the peak estradiol was 5757 pmol/ml. Five follicles were documented by ultrasound, measuring 18–20 mm average diameter, and totally twelve follicles were recorded. Oocyte retrieval was performed 36 hours after recombinant human chorionic gonadotropin (r-hCG,Ovidrel^®^,Merck Serono, Germany) administration. No recognizable oocytes were retrieved after thorough aspiration and flushing of the seven follicles in the right ovary, so the oocyte retrieval was suspended and the HCG level in blood was evaluated immediately. Because of the normal hormones assays, ovum pick up (OPU) was performed 2 hours later, and five follicles in the left ovary were aspirated. Several cumulus–corona complexes were recovered but no recognizable oocytes inside. The second IVF attempt was carried out with GnRH antagonist protocol (Table S3) 3 months later; dual trigger was administered. However, similar to the first attempt, no recognizable oocytes were retrieved.

### Case 2

The sister of the proband (Figure 2A II-2) was 28 years old and suffered from primary infertility for five years. The semen analysis of her husband presented mild asthenozoospermia. This couple underwent IVF treatment at the same center and by the same team half a year later. GnRH antagonist protocol (Table S3) was adopted, and dual trigger was administrated. Nine cumulus–corona complexes were collected (Figure 1B), however, no recognizable oocytes were found as well. The cumulus–corona complexes were digested by hyaluronidase (HYASE^TM^-10X, Vitrolife, Sweden).

### Genetic analysis

All genomic DNA from all six family members were extracted from the peripheral blood by Qiagen QIAamp DNA blood mini kit according to the manufacturer’s instructions. To determine the causative mutation, we carried out Whole-exome sequencing (WES) on the proband of the family (Figure 2A II-3). Sequencing library was constructed using Agilent SureSelect V6 kit according to the manufacturer’s manual. Raw sequencing reads, i.e., FASTQ files, were produced by Hiseq4000 sequencer (Illumina). Raw sequencing reads were aligned to the human reference genome (hg19) and variants in forms of SNV, indel, CNV, and SV were called by MERAP[21]. Variants were first filtered against 1000 Genome, ExAc database, ESP6500 database, and an in-house ethnic-matched 2,200 individual genomes, in order to remove those with MAF > 0.01. Candidate genes were then assigned to those with variants predicted damaging by SIFT, MutationTaster2, and PolyPhen2 in consensus and in evolutionarily conserved regions defined by PhyloP and GERP. Annotation of candidate variants was based on RefSeq gene models. After evaluating clinical relevance of each gene by OMIM and HGMD, the candidate variants were collected (Table S5).

The cosegregation test was conducted for the selected candidate variants on all six family members. First, PCR amplification was performed using Q5 High-Fidelity Polymerase (5x, NEB, MA, USA) with the specific primers targeted to the selected candidate variants (Table S6). Then, the PCR products were illustrated by the Sanger Sequencing to determine the carrier status of the variants.

## Discussion

Empty follicle syndrome is defined failure to retrieve oocytes during *in vitro* fertilization (IVF) protocols. It was classified as false and genuine. “False” EFS is mainly caused by pharmacological or iatrogenic problems, it due to incorrect use human chorionic gonadotrophin (HCG) in IVF treatment for oocyte maturation. So it can be rescured by corresponding correction measures, such as change another batch of HCG administration[5–7], employed recombinant HCG[8], endogenous gonadotrophin surge induced by GnRH agonist in an antagonist cycle[9], and dual trigger for oocyte maturation[10]. Patients in the afore-mentioned practices responded well to the respective rescue protocols, and a relapse could be avoided granted the appropriate treatment. However, recurrent EFS were reported by different teams after applying these improved measures [11, 12]. For example, a case report presented two sisters who underwent three cycles of controlled ovarian hyper-stimulation with normal follicular development, estradiol levels and optimal b-hCG levels, but without oocytes or cumulus-corona complexes retrieved[13]. Such cases implicate that refractory “genuine” EFS do exist, which may well have underlying genetic defects.

A pericentric inversion of chromosome 2, namely, “46, XX, inv(2)(p11q21)” was identified by karyotyping a patient who had multiple failed IVF attempts, due to the absence of oocyte and granulosa cells in the follicular fluid. The authors hypothesized that chromosomal abnormality was ascribed to the etiology of EFS [22], although the evidence was circumstantial. *LHCGR* gene Mutations were definitely etiologies of EFS, which were confirmed by different teams [14, 15], *LHCGR* gene encodes the luteinizing hormone/choriogonadotropin receptor, the impaired functioning of LHCGR caused diminished response to the administration of beta-hCG. The correlation of *LHCGR* mutation with EFS was substantiated in female mice lacking *Lhcgr*, which had no pre-ovulatory follicles or corpora luteal, likely secondary to defects in follicular maturation and ovulation, analogous to the human EFS phenotype [18].

Gene mutation in zona pellucida glycoprotein is another definitely aspect of etiologies. Zona pellucida glycoprotein 3 (ZP3) was identificated in two EFS families [16]. The author proposed that via dominant negative inhibition, this mutation could destroy the assembly of the zona pellucid (ZP) thus leading to oocyte degeneration and EFS.

In this study, we identified compound heterozygous mutations of *ZP1* gene from a family with two sisters of abnormal oogenesis with EFS. The allelic mutants were transmitted from either parent. We hypothesized that these mutations hurdled the maturation of oocyte at an early developmental stage, incurring degeneration and empty follicles.

The model of zona pellucida development includes intraplasmic synthesis and maturation of the component proteins, transportation to the cell surface, and interweaving among the four ZP proteins [20]. Among this process, ZP1 plays a critical role which cross-links the long chains of filaments made of consecutive alternates of ZP2, ZP3, and ZP4. This model renders a convincing explanation why ZP1 is indispensable in formation of zona pellucida while the expression of ZP1 is relatively lower than the three other components.

In Rankin et al.’s work in 1999[18], the oocyte zona pellucida of the mouse line (Zp1-/-) was more loosely organized than those around normal oocytes. The ovulated eggs were comparable with the control ones in numbers, while the subsequent fertilization was suboptimal, causing litters significantly smaller than those produced by normal mice. This work substantiated the indispensable role of ZP1 in oocyte maturation, either in human or mice.

We noticed that in Huang’s work, the frameshift homozygote in *ZP1* truncated the external hydrophobic patch, the consensus furin cleavage site, the transmembrane domain, the cytoplasmic tail, and the second half of the zona pellucida domain, while the N-terminal signal sequence, the trefoil domain, and the first half of the zona pellucida domain were remained[20]. While in our case, the more severe condition, namely, without observable oocyte, could be explained by the remaining half amount of the previously reported truncated ZP1 protein, which was insufficient to maintain the oocyte normal development, so the maturation of oocyte was hurdled at an early developmental stage leading to degeneration and empty follicles.

## Conclusion

In our work, we proposed a scenario in which allelic mutations in the *ZP1* gene may well induce refractory “genuine” EFS, so besides the recessive EFS caused by *LHCGR* gene mutations and the dominant EFS caused by *ZP3* gene mutations, we added another candidate into the EFS genetic repertoire, providing a novel glimpse into the heterogeneity of EFS etiology.

## Authors’ roles

Study was conceived by L.S. and N.L.; Z-H.C. and H-W.Z. C-L. L was in charge of blood samples collected of the family. L.S., Y.D. and M-N.Y. performed IVF treatment and collected the clinic data. Z-H.C conducted cumulus–corona complexes collection. N.L. conducted the genetics analysis. X.F. performed in silico analyses. X-F.P., T.X. and Q-Y.Z. performed the cosegragation study. L.S., H-W Z., Z-H.C. and N.L. were in charge of the revisions of the paper. All authors approved the final manuscript.

---

**Table S1.** Clinic characteristics:

|  | Proband (II-3) | Sister(II-2) |
| --- | --- | --- |
| Age (y) | 23 | 28 |
| BMI | 21.1 | 16.9 |
| Menarche (y) | 16 | 14 |
| menses length (d) | 30 | 28 |
| Infertility duration (y) | 3 | 5 |
| Type of infertility | Primary infertility | Primary infertility |
| Ovarian reserve |  |  |
| Antrol follicles | 12 | 9 |
| Basal FSH (IU/L) | 6.32 | 4.94 |
| Basal LH (IU/L) | 2.69 | 3.07 |
| Basal E2 (pmol/L) | 83 | 136 |
| AMH (ng/ml) | 5.64 | 6.1 |

**Table S2.**
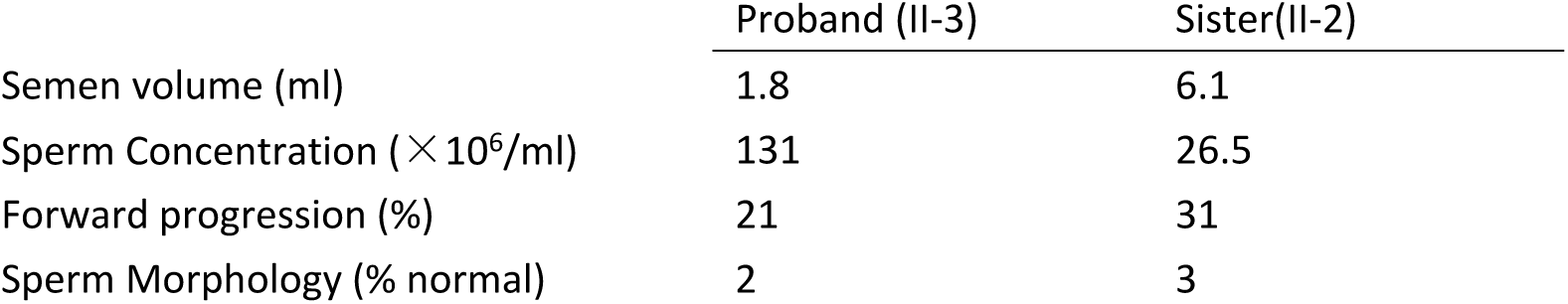
Semen analysis:

**Table S3.**
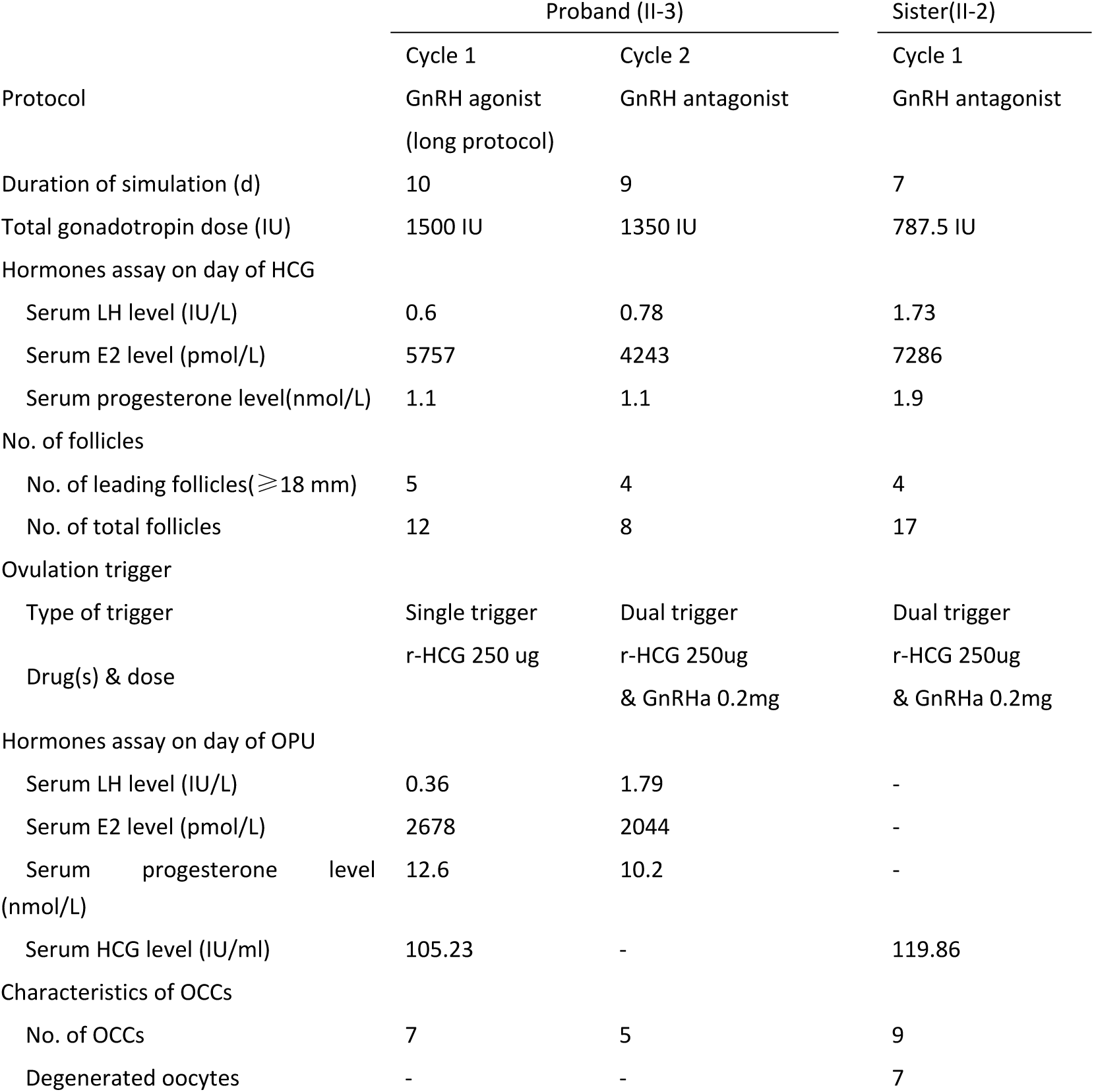
The ART cycles description.

**Table S4.** Coding-coverage of the Whole-exome sequencing result for the proband of the family:

|  | Coding-coverage(%) |
| --- | --- |
| 10X | 97.85 |
| 20X | 96.29 |
| 30X | 92.38 |
| 40X | 86.03 |
| 50X | 78.31 |

**Table S5.** The candidate mutations of *ZP1* gene:

| Gene name | Mutations |
| --- | --- |
| ZP1(NM_207341,zona pellucida glycoprotein 1 (sperm receptor),638aa) | g.11:60640688_60640695del,c.1166_1173del,p.389fs |
| ZP1(NM_207341,zona pellucida glycoprotein 1 (sperm receptor),638aa) | g.11:60635195_60635199del,c.161_165del,p.54fs |

**Table S6.**
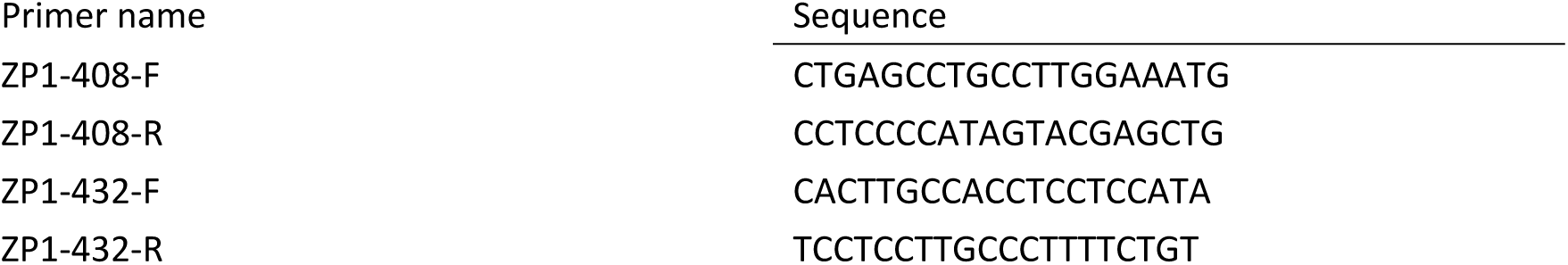
Primer for PCR confirmation:

Dr. Ling Sun gained her MD in 1995 and her PhD in 2007. She is a gynecologist, active in the fields of reproductive medicine; she set up the Department of Assisted Reproduction in Guangzhou women and children’s hospital in 2009. Her main research interests are the genetics of infertility.

